# Mechanical Disturbance of Trichomes Enhances Cd^2+^ Uptake via Malate-Mediated Stomatal Opening in *Arabidopsis*

**DOI:** 10.1101/2025.11.20.689625

**Authors:** Juan Du, Hongyang Wei, Tongxin Shen, Xue Chang, Mingyu Hao, Yang Qi, Ning Liu, Zhiyan Cao, Jingao Dong, Li Hong Zhou

## Abstract

Phytoremediation of cadmium-contaminated soils relies on efficient metal uptake and translocation in plants. We previously found that gentle mechanical stimulation of non-glandular trichomes by brushing significantly enhances shoot Cd²⁺ accumulation in *Arabidopsis thaliana*, although the underlying physiological mechanism remained unclear. Here, we demonstrate that trichome brushing specifically induces stomatal opening, elevates transpiration, and promotes Cd²⁺ uptake, as evidenced by measurements of stomatal aperture, stomatal conductance, transpiration rate, and Cd²⁺ concentration in wild-type and mutant leaves. These responses are absent in the glabrous *gl1-1* mutant and are not elicited by brushing epidermal pavement cells, indicating a trichome-specific mechanosensory pathway. Mechanistically, this pathway requires MPK3/MPK6 activity and employs malic acid as a key regulator of stomatal aperture, functioning independently of canonical ABA signaling. Our findings uncover a previously unrecognized trichome-to-stomatal signaling cascade that links trichome mechanostimulation to enhanced heavy metal accumulation.

## Introduction

Cadmium (Cd²⁺) is a highly toxic heavy metal with a biological half-life of 10 to 30 years in humans. Due to its severe health risks, the World Health Organization and the United Nations Environment Programme identified Cd²⁺ as a priority food contaminant in 1972 (Wagner, 1993). In China, anthropogenic activities—particularly industrial emissions and certain agricultural practices—have contributed to widespread Cd²⁺ pollution in cultivated soils (Wen et al., 2022). Grain contamination is estimated to reach 13 million tonnes annually (Wang et al., 2019), presenting a significant threat to human health through the food chain. As a result, effective remediation of Cd²⁺-contaminated soils has become an urgent environmental and public health priority.

Heavy metals such as Cd²⁺ are resistant to degradation, creating persistent environmental challenges. Conventional remediation strategies, including phyto-stabilisation (Shackira & Puthur, 2019) and chemical passivation (Lan et al., 2021), can reduce metal bioavailability but do not remove contaminants from the soil (Bolan et al., 2011). In contrast, phytoextraction employs hyperaccumulating or high-biomass plants to absorb metals and translocate them to harvestable shoots (Suman et al., 2018). Processing this biomass enables potential metal recovery, offering a sustainable route to soil decontamination (Ghori et al., 2016). Efficient phytoextraction depends on maximizing heavy metal accumulation in aerial tissues. While most research has focused on genetic regulation of metal transport and tolerance (Pasricha et al., 2021), the contribution of plant structural features—such as epidermal trichomes—remains underexplored.

Our previous work showed that *A. thaliana* trichomes possess a mechanically sensitive architecture (Liu et al., 2016; Zhou et al., 2017). Inducing this instability through gentle mechanical stimulation—such as brushing, insect contact, or acoustic resonance—significantly increases shoot Cd²⁺ accumulation (Guo et al., 2022; Liu et al., 2017; Peng et al., 2022). However, the physiological and molecular mechanisms linking trichome mechanostimulation to enhanced Cd²⁺ accumulation remain unknown.

Plants integrate multiple signalling networks to respond to environmental cues, including heavy metals and mechanical stimuli. Candidate pathways include abscisic acid (ABA) signalling, mitogen-activated protein kinase (MAPK) cascades, calcium (Ca²⁺) signalling, reactive oxygen species (ROS) production, and microRNA regulation (Shen et al., 2022; Opdenakker et al., 2012; Kaur et al., 2021; Romero-Puertas et al., 2019; Ding et al., 2018; Lu et al., 2024). Cd²⁺ exposure can activate MPK3 and MPK6 via ROS, leading to downstream responses such as phytochelatin synthesis (Liu et al., 2010; Chen et al., 2016). ABA signalling generally acts to limit Cd²⁺ uptake (Pan et al., 2020; You et al., 2022). Mechanical stimulation of trichomes is also known to elicit intercellular Ca²⁺ waves (Matsumura et al., 2022) and involves regulators such as CAMTA3 in touch responses (Darwish et al., 2022). Yet, it remains unclear which of these pathways, if any, transmit trichome-derived mechanical signals to modulate Cd²⁺ accumulation.

Here, we address this knowledge gap by investigating the physiological and signalling events linking trichome mechanostimulation to shoot Cd²⁺ accumulation in *A. thaliana*. We investigated how trichome brushing affects stomatal behavior—including aperture, conductance, and transpiration—and assessed the involvement of CAMTA3, ABA and MAPK signaling pathways. Our results demonstrate that trichome stimulation activates a MAPK-dependent pathway leading to malate-mediated stomatal opening and increased Cd²⁺ uptake, revealing a previously uncharacterized mechanosensory cascade from epidermal trichomes to stomatal regulation

## Materials and methods

### 3.1 Plant growth

*A. thaliana* wild type (WT, Col 0), mutants (*gl1-1*, *sr1-2*, *mpk3-1*, *mpk6-4*), and transgenic (*SR1 ox5*) lines were used in this study

For soil-grown plants, seeds were sown in trays containing Pindstrup Plus Blue peat substrate soil (Pindstrup, Denmark) and vernalized at 4 °C for 2 days. The trays were then transferred to a controlled environment growth room (22 °C, 40–70% relative humidity, 16 h light/8 h dark photoperiod) and watered daily with 500 mL of tap water. After 2–3 weeks of growth, plants received a one-time application of 1500 mL of 5 µM CdCl₂ solution, which was fully absorbed by the soil. Daily watering (500 mL) was resumed for an additional 3–7 days before sampling.

For plate-grown plants, seeds were surface sterilized in 75% ethanol for 4 min, followed by 100% ethanol for 1 min, and then rinsed 2–3 times with sterile water. Sterilized seeds were evenly spotted on half-strength Murashige and Skoog (½ MS) agar plates, with or without 1 µM CdCl₂. Plates were vernalized at 4 °C for 2 days, then placed vertically in a growth room (22 °C, 40% relative humidity, 16 h light/8 h dark) for 2–4 weeks.

### 3.2 Cd²⁺ Quantification in Leaf Tissue

WT and mutant *A. thaliana* plants were grown on ½ MS plates for 2–4 weeks. Fully expanded leaves were subjected to gentle mechanical stimulation by brushing trichomes 20 times with a 3 mm tip brush, then left to rest for 2 h. Fresh weight was recorded prior to mineralization. The leaves were immersed in 1 mL of HNO₃ for 12 h, followed by digestion at 110°C for 2.5 h. The digests were then diluted with pure water (Hangzhou Wahaha Group Co., Ltd.) to a final volume of 4.5 mL. A 1 mL aliquot was further diluted 10-fold with pure water in a 10 mL centrifuge tube and passed through a 0.22 µm aqueous filter into a fresh tube. Cd²⁺ concentrations in the filtrates were determined by inductively coupled plasma mass spectrometry (ICP-MS; NexION 300X, PerkinElmer, USA).

### 3.3 Measurements of Stomatal Aperture, Stomatal Conductance, and Transpiration Rate

WT, mutant, and transgenic plants were grown on MS plates. Trichomes were mechanically stimulated as described in Section 3.2, and plants were rested for 2 h before measurement. The first two true leaves or upper epidermal peels from two leaves per plant were examined under a light microscope (Olympus BX53F). For each plant, 50 stomata were imaged, and stomatal aperture was quantified as the width-to-length ratio using ImageJ software.

For stomatal closure assays, 20 µL of 100 nM ABA solution was applied to the leaf surface for 1 h. To assess the effect of malic acid, detached leaves from WT or *gl1-1* plants were floated on stomatal buffer solution (10 mM MES, 50 mM KCl, 10 mM CaCl₂, pH 6.5) with or without 0.1 mM malic acid for 3 h. For malic acid dehydrogenase (MDH) treatment, detached leaves were floated on 0.1 M MES (pH 7.5) containing 0.1 mM MDH in the dark for 3 h. After treatment, upper epidermal peels were prepared, and stomatal apertures were measured as described above.

For soil-grown plants (4 weeks old), fully expanded leaves from WT, *gl1-1*, *mpk3-1*, and *mpk6-4* genotypes were used to measure stomatal conductance and transpiration. Following gentle brushing of trichomes or epidermal pavement cells, plants were allowed to recover for 2 h. Two similarly sized leaves (typically the fifth or sixth true leaf pair) were selected per plant, and measurements were taken using a portable photosynthesis system (LI-6800; LI-COR Biosciences, USA).

### 3.4 Scanning Electron Microscopy (SEM) Imaging

Fully expanded leaves from 10–14-day-old WT and *gl1-1 Arabidopsis* plants grown on MS agar plates were brushed 20 times with a 3 mm-tip brush to stimulate trichomes and epidermal pavement cells, then allowed to rest for 2 h. The first two true leaves were excised and fixed in 2.5% glutaraldehyde. Fixed samples were centri-fuged at 4000 *rpm* for 5 min to remove excess fixative, then dehydrated through a graded ethanol series (30%, 50%, 70%, 90%, and 100%), with each step lasting 15–20 min. After dehydration, samples were wrapped and transferred to a critical point dryer (Tousimis Autosamdri-815). Dried leaves were mounted on aluminium SEM stubs and sputter-coated with gold using a vacuum sputter coater (Hitachi MC1000).

SEM imaging was performed using a field-emission scanning electron microscope (FE-SEM; SU8020, Hitachi High-Tech, Japan) operated at an accelerating voltage of 5 kV. Images were captured using a secondary electron detector at magnifications ranging from 200× to 500×. All samples were imaged under high vacuum conditions, and representative micrographs were selected from at least three biological replicates.

### 3.5 Dithizone Staining of Cd²⁺ in Seedlings

*A. thaliana* seedlings (10–20 days old) grown on ½ MS plates containing varying concentrations of Cd²⁺ were stained with dithizone (Sigma, USA) following the method of Seregin and Ivanov (1997). The staining solution was prepared by dissolving 0.0052 g dithizone in 10 mL of acetone, adding 3 mL of distilled water, and then supplementing with 100 µL of glacial acetic acid. Seedlings were immersed in this solution for 4 h, then transferred to 50% (v/v) ethanol for 1 h to bleach chlorophyll. After bleaching, plants were rinsed in sterile water for 30 min. Samples were observed directly under a light microscope (Olympus BX53F) to detect red precipitates formed by the reaction of dithizone with Cd²⁺ ions.

### 3.6 LC–MS Detection of ABA and Malic Acid

WT and *gl1-1* mutant *A. thaliana* plants were grown on MS plates and subjected to mechanical stimulation as described in Section 3.2. Leaf samples were immediately frozen in liquid nitrogen and ground to a fine powder. Metabolites were extracted by suspending the powder in 0.1% acetic acid in acetonitrile and incubating at 4 °C overnight. Extracts were centrifuged at 12 000 *rpm* for 1 min, and 1 mL of the supernatant was mixed with 10 mg C18 powder for purification. The mixture was centrifuged at 4 000 *rpm* for 5 min, and the resulting supernatant was filtered through a 0.22 µm membrane to obtain the final extract for analysis.

Chromatographic separation was performed on a Kinetex 2.6 µm C18 100 Å reversed-phase column. The mobile phase consisted of 0.1% formic acid in water (Phase A) and acetonitrile (Phase B). The elution program was: 0–1 min, 10% B; 1–min, linear gradient from 10% to 90% B; 4–5 min, 90% B; 5–8 min, return to 3% B. The flow rate was 0.25 mL min⁻¹, column temperature 40 °C, and sample tray temperature 23 °C. Injection volume was 5 µL, with a total run time of 8 min.

Mass spectrometric detection was carried out using an electrospray ionization (ESI) source at 4 500 V in multiple reaction monitoring (MRM) mode. Curtain gas pressure was 30 psi, ionization voltage 9 V, source temperature 450 °C, spray gas pressure 65 psi, and auxiliary heating gas pressure 50 psi.

### 3.7 Quantitative Real-Time PCR (qRT-PCR) Analysis

Appropriate amounts of *A. thaliana* leaves were frozen in liquid nitrogen and ground to a fine powder. Total RNA was extracted using the Plant Extractable RNA Kit (UNIQ-10 Column Trizol Total RNA Isolation Kit; Bioengineer, China) following the manufacturer’s instructions. Two micrograms of total RNA were reverse-transcribed into cDNA. Quantitative real-time PCR was performed with gene-specific primers using AugeGreen qPCR Master Mix (TaKaRa, RR820A). *ACTIN2* (At3g18780) was used as the internal reference gene. Relative transcript levels were calculated using the comparative Ct method (2⁻ΔΔCt), with expression values normalized to those of the wild-type (WT) control.

### 3.8 Statistics

All value data reported in this paper are expressed as mean ± standard deviation. Means were compared using Student’s *t-test*, with a threshold of p < 0.05 for statistical significance. Each experiment was repeated at least three times with independent biological replicates. For each biological replicate, multiple technical replicates (e.g., leaves or stomata per plant) were averaged before statistical analysis. Statistical significance: ns, not significant (p > 0.05); **, p < 0.01;***, p < 0.001; ****, p<0.0001.

## Results

### 4.1 Stimulation of trichomes, not epidermal pavement cells, enhances Cd²⁺ accumulation in *A. thaliana* shoots

Our previous work demonstrated that mechanical brushing of trichomes significantly increases Cd²⁺ concentration in the shoots of *A. thaliana* (Guo et al., 2022). Because brushing trichomes with a paintbrush inevitably involves contact with adjacent epidermal pavement cells, we asked whether stimulation of pavement cells alone contributes to Cd²⁺ uptake. To address this, we compared trichome brushing in WT plants with pavement cell brushing in the glabrous mutant *gl1-1*.

Consistent with earlier findings, trichome brushing in WT plants markedly increased the presence of large red Cd²⁺–dithizone precipitates within trichomes (Fig. 1b), whereas such precipitates were rarely observed prior to brushing (Fig. 1a). ICP-MS analysis confirmed a significant rise in shoot Cd²⁺ concentration in WT plants, both under Cd²⁺ stress (Fig. 1c) and in its absence (Fig. S1c). In contrast, brushing the epidermal pavement cells of *gl1-1* leaves had no detectable effect on Cd²⁺ accumulation (Fig. 1d, e), with ICP-MS measurements showing no significant change in shoot Cd²⁺ concentration regardless of Cd²⁺ exposure (Fig. 1f, Fig. S1f).

**Fig. 1.**
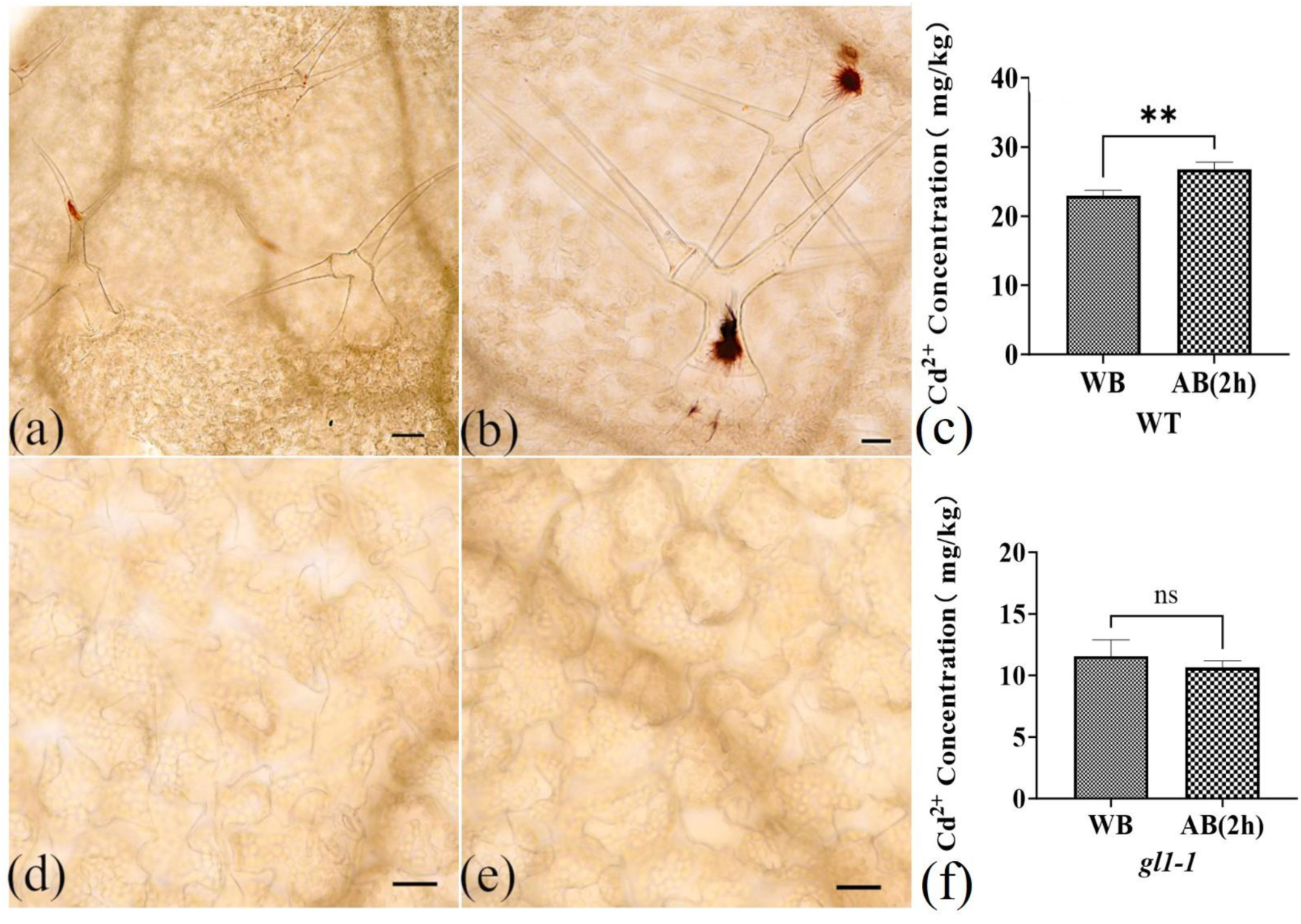
Trichome brushing, but not epidermal pavement cell brushing, enhances Cd²⁺ accumulation in *A. thaliana* shoots. (a, b) Dithizone staining of WT leaves before (a) and after (b) 2 h of targeted trichome brushing shows intensified Cd^2+^–dithizone precipitates (red or dark red) within trichomes following stimulation. (d, e) *gl1-1* mutant leaves stained before (d) and after (e) epidermal brushing exhibit no detectable change in Cd²⁺ localization, indicating trichome-specific responsiveness. (c, f) Shoot Cd²⁺ concentrations quantified by ICP-MS in WT (c) and *gl1-1* (f) plants grown on Cd²⁺-supplemented MS plates. WB, without brushing; AB (2 h), after 2 h of brushing. Scale bar = 100 µm. Statistical significance: ns, not significant (p > 0.05); **, p < 0.01.

Together, histochemical staining and ICP-MS quantification demonstrate that mechanical stimulation of trichomes — but not epidermal pavement cell contact — promotes Cd²⁺ accumulation in *A. thaliana* shoots.

### 4.2 Stimulating trichomes enhances stomatal aperture, conductance, and leaf transpiration

Leaf trichomes are known to reduce transpiration and lower leaf temperature by increasing boundary layer resistance (Bickford, 2016). Because transpiration dynamics strongly influence Cd²⁺ accumulation within trichomes and its distribution across aerial tissues (Salt et al., 1995), we investigated whether mechanical destabilization of trichomes alters leaf transpiration by evaluating its impact on stomatal aperture.

Mechanical brushing of WT trichomes significantly increased stomatal aperture (Fig. 2a–c; Fig. S2). This effect was also observed in plants grown without Cd²⁺ supplementation (Fig. S3a–c), demonstrating that the response is independent of Cd²⁺ stress. In contrast, brushing the epidermal surface of *gl1-1* mutant leaves did not affect stomatal aperture under either control or Cd²⁺ conditions (Fig. 2d–f; Fig. S3d–f).

**Fig. 2.**
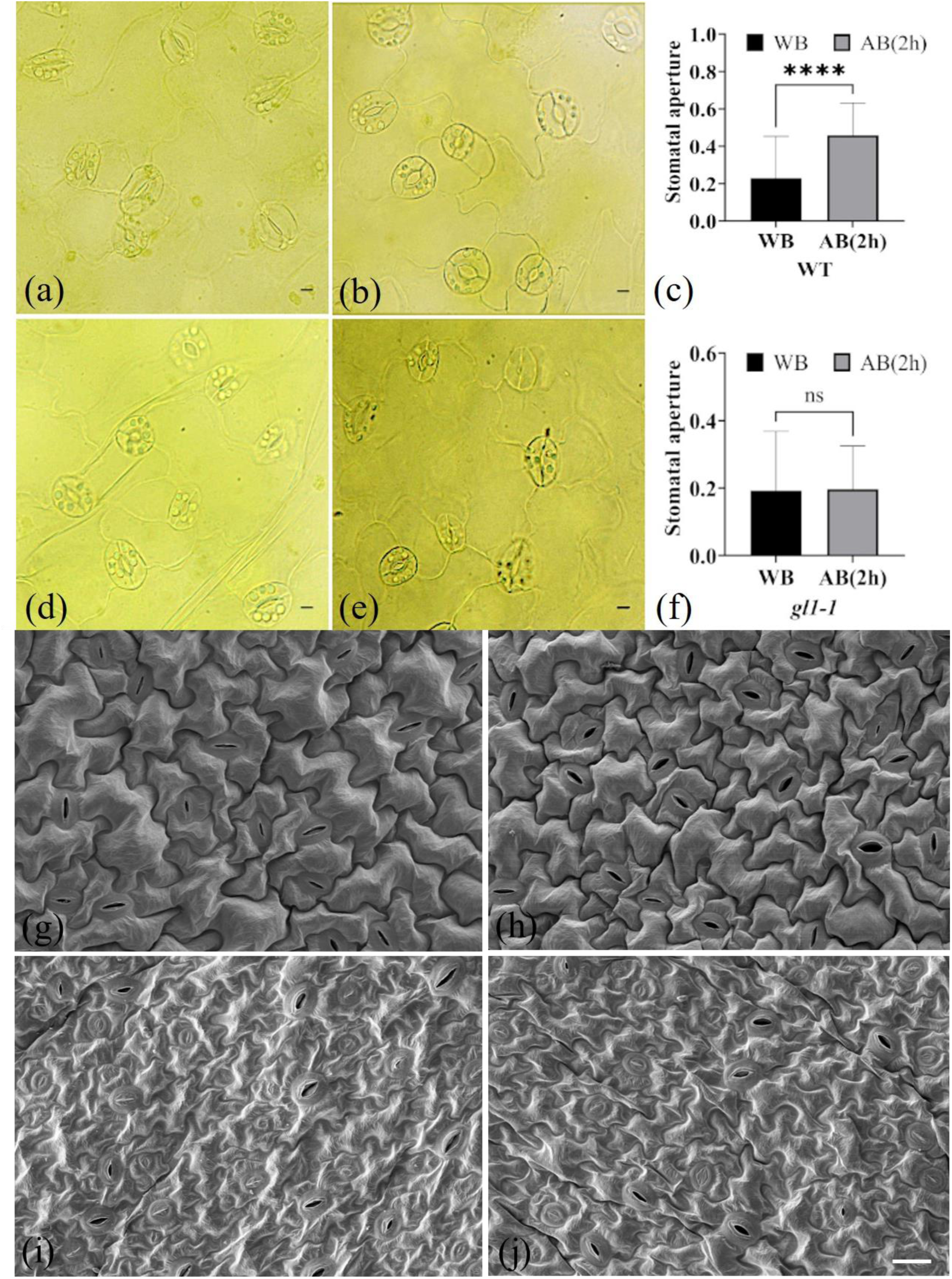
Mechanical stimulation of trichomes promotes stomatal opening in WT *Arabidopsis* leaves. (a, b) Representative stomata in WT leaves without (a) and with (b) trichome brushing. (d, e) Representative stomata in *gl1-1* mutant leaves without (d) and with (e) epidermal brushing. (c, f) Quantification of stomatal aperture in WT (c) and *gl1-1* (f) leaves following brushing treatments. (g, h) Representative SEM images of WT leaves without (g) and with (h) trichome brushing. (i, j) Representative SEM images of *gl1-1* mutant leaves without (i) and with (j) epidermal brushing. Plants in panels (a–f) were grown on Cd²⁺-supplemented MS plates; those in (1–j) were cultivated on Cd²⁺-free MS plates. Scale bars: 40 µm (a–f); 20 µm (g–j). Statistical significance: ns, not significant (p > 0.05); ****, p < 0.0001.

To verify the morphological consequences of trichome brushing, we examined leaf surfaces using SEM. Comparisons of brushed and unbrushed WT and *gl1-1* leaves confirmed that trichome brushing selectively increased stomatal aperture in WT (Fig. 2g–h), whereas epidermal brushing of *gl1-1* leaves had no detectable effect (Fig. 2i-j). Together, these results demonstrate that mechanical perturbation of trichomes specifically triggers stomatal opening.

Changes in stomatal aperture directly influence transpiration rate and stomatal conductance. To determine whether trichome stimulation modulates these physiological parameters, we performed gas exchange measurements using the Li-6800 Portable Photosynthesis System. Brushing trichomes in WT leaves significantly increased both transpiration rate and stomatal conductance (Fig. 3; Fig. S4). In contrast, brushing the epidermal surface of *gl1-1* mutant leaves had no measurable effect on either trait (Fig. 3). These results suggest that mechanical stimulation of trichomes enhances Cd²⁺ accumulation by promoting stomatal opening and thereby increasing transpiration-driven uptake.

**Fig. 3.**
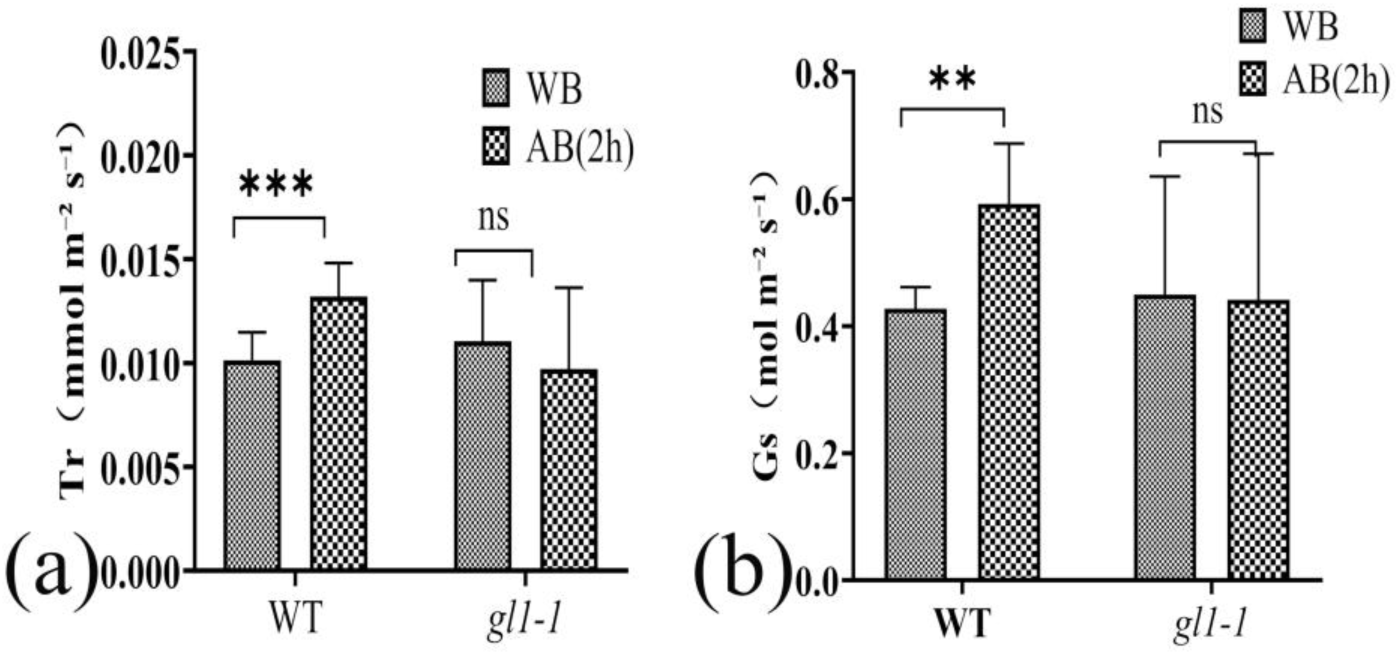
Trichome stimulation enhances transpiration rate and stomatal conductance in WT *Arabidopsis* leaves. (a, b) Quantification of (a) transpiration rate and (b) stomatal conductance in WT and *gl1-1* mutant leaves with or without mechanical stimulation via brushing. Each condition includes six biological replicates. All plants were grown in soil supplemented with CdCl₂.Statistical significance: ns, not significant (p > 0.05); **, p < 0.01, ***p < 0.001

### 4.3 Stomatal opening induced by trichome stimulation is independent of ABA signaling

Given the pivotal role of ABA in regulating stomatal aperture and mediating responses to Cd²⁺ stress, we investigated whether trichome-induced stomatal opening is linked to the ABA signaling pathway. ABA concentrations were quantified in WT and *gl1-1* leaves before and after brushing. In WT leaves, ABA levels increased from 0.02 ± 0.0012 µg/g to 0.06 ± 0.0002 µg/g following trichome stimulation. In contrast, *gl1-1* leaves exhibited a decrease in ABA concentration from 0.04 ± 0.0021 µg/g to 0.03 ± 0.0004 µg/g (Fig. 4a). These results indicate that trichome brushing elevates ABA concentration in WT leaves, whereas epidermal brushing in trichome-deficient *gl1-1* reduces ABA accumulation.

**Fig. 4.**
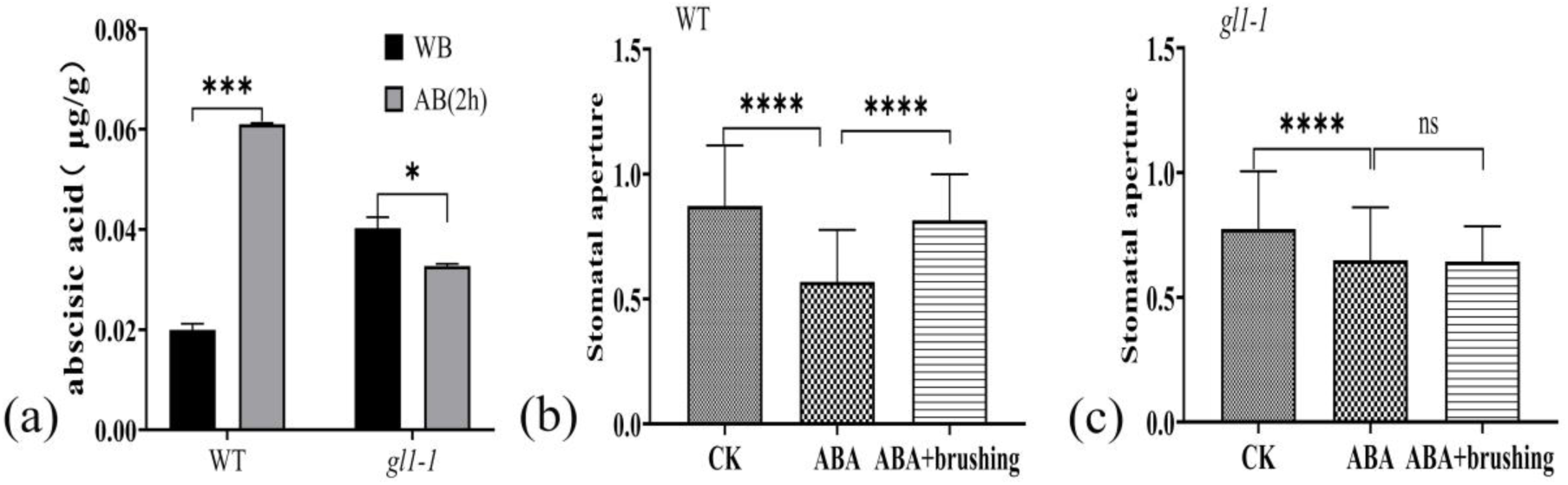
Brushing trichomes increases stomatal aperture despite exogenous ABA application. (a) ABA concentrations in WT and *gl1-1* leaves before brushing (WB, without brushing) and 2 h after trichome stimulation (AB 2h, after brushing). (b, c) Stomatal aperture measurements in WT (b) and *gl1-1* (c) leaves following ABA treatment and subsequent brushing. For ABA treatment, fully expanded leaves were incubated in 100 nM ABA solution for 1 h. The solution was then gently removed, and leaves were brushed 20 times with a 3 mm-tip brush, followed by a 2 h resting period before stomatal aperture measurement. All plants were cultivated on Cd²⁺-supplemented MS medium. Statistical significance: ns, not significant (p > 0.05); ***, p < 0.001;****, p < 0.001.

Typically, elevated ABA levels are associated with stomatal closure. However, our data reveal a paradoxical increase in stomatal aperture following trichome brushing, suggesting that this response occurs independently of canonical ABA signaling. To further test this hypothesis, we applied 100 nM ABA to the leaf surfaces of WT and *gl1-1* plants. As expected, ABA treatment induced widespread stomatal closure (Fig. S5). Remarkably, subsequent brushing of WT trichomes still promoted stomatal opening despite prior ABA application (Fig. 4b), whereas *gl1-1* leaves showed no significant change in aperture (Fig. 4c). Similar results were obtained in plants grown on MS medium without Cd²⁺ supplementation (Fig. S6), confirming that the effect is not contingent on metal stress.

Collectively, these findings demonstrate that mechanical stimulation of trichomes overrides ABA-induced stomatal closure and promotes aperture expansion through an ABA-independent pathway.

### 4.4 Stomatal opening induced by trichome stimulation is independent of CAMTA3

Mechanical stimulation of trichomes is known to initiate intercellular calcium waves and activate CAMTA3-dependent transcriptional programs involved in biotic stress responses and touch-induced gene expression (Matsumura et al., 2022; Darwish et al., 2022). To determine whether CAMTA3 contributes to Cd²⁺ accumulation or stomatal regulation, we analyzed stomatal aperture changes and Cd²⁺–dithizone precipitate formation in *sr1-2* (*CAMTA3* loss-of-function mutant) and *SR1-ox5* (*CAMTA3*-overexpressing) lines before and after trichome brushing.

Under Cd²⁺ stress conditions, trichome stimulation significantly increased stomatal aperture in both genotypes (Fig. 5f–h; 5k–m), and similar responses were observed in Cd²⁺-free conditions (Fig. S7). In *sr1-2* mutants, stomatal aperture increased following trichome brushing (Fig. 5h), and *SR1-ox5* plants showed a comparable increase (Fig. 5m). Additionally, Cd²⁺ accumulation—visualized as red dithizone precipitates—was detected in trichomes of both *sr1-2* and *SR1-ox5* plants, with enhanced deposition after brushing (Fig. 5d, 5e; 5i, 5j; 5n, 5o).

**Fig. 5.**
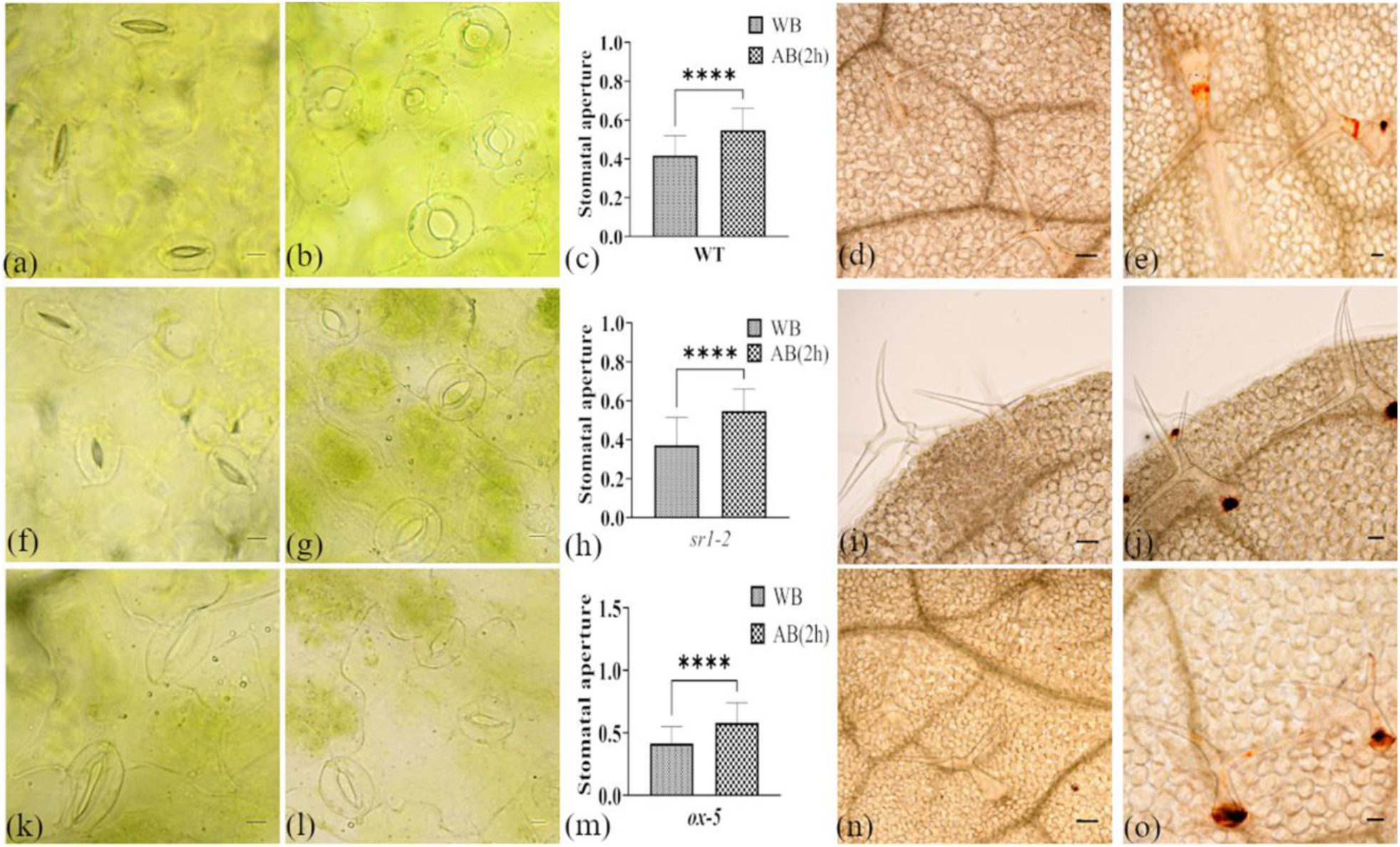
Trichome stimulation increases stomatal aperture and Cd²⁺ accumulation in WT and *CAMTA3*-modified *Arabidopsis* lines. Representative stomata and histological analysis of Cd²⁺ localization in unstimulated (a, f, k; d, i, n) and mechanically brushed (b, g, l; e, j, o) leaves of WT (a, b; d, e), *sr1-2* (f, g; i, j), and *SR1-ox5* (k, l; n, o) plants grown on Cd²⁺-supplemented MS plates. Stomatal aperture was quantified in epidermal peels from WT (c), *sr1-2* (h), and *SR1-ox5.* Statistical significance: ****, p < 0.0001.

These findings demonstrate that both *CAMTA3*-deficient and *CAMTA3*-overexpressing lines exhibit stomatal aperture responses and Cd²⁺ accumulation patterns comparable to WT plants. We conclude that CAMTA3 is not required for trichome-destabilization-induced stomatal opening or Cd²⁺ sequestration, indicating that these processes operate independently of CAMTA3-mediated signaling.

### 4.5 Trichome-stimulation-induced stomatal opening requires *MPK3* and *MPK6*

Mechanical stimulation of trichomes activates phosphorylation of MPK3 and MPK6, key components of the MAPK cascade that orchestrates biotic stress responses and touch-induced gene expression (Matsumura et al., 2022). These kinases are also known to regulate stomatal closure during pathogen defense (Su et al., 2017). Given that neither ABA signaling nor CAMTA3 mediates the trichome-induced increase in stomatal aperture, we hypothesized that MAPK signaling may serve as a critical regulator of stomatal dynamics and Cd²⁺ accumulation in response to mechanical stimulation.

To test this, we examined stomatal aperture changes in *mpk3-1* and *mpk6-4* mutants before and after trichome brushing. Under both Cd²⁺-exposed (Fig. 6) and Cd²⁺-free conditions (Fig. S8), trichome stimulation failed to induce stomatal opening in either mutant. Quantitative measurements revealed no statistically significant differences in stomatal aperture before and after brushing (Fig. 6f, 6i; Fig. S8f, S8i), indicating that *MPK3* and *MPK6* are essential for the trichome-mediated stomatal opening response.

**Fig. 6.**
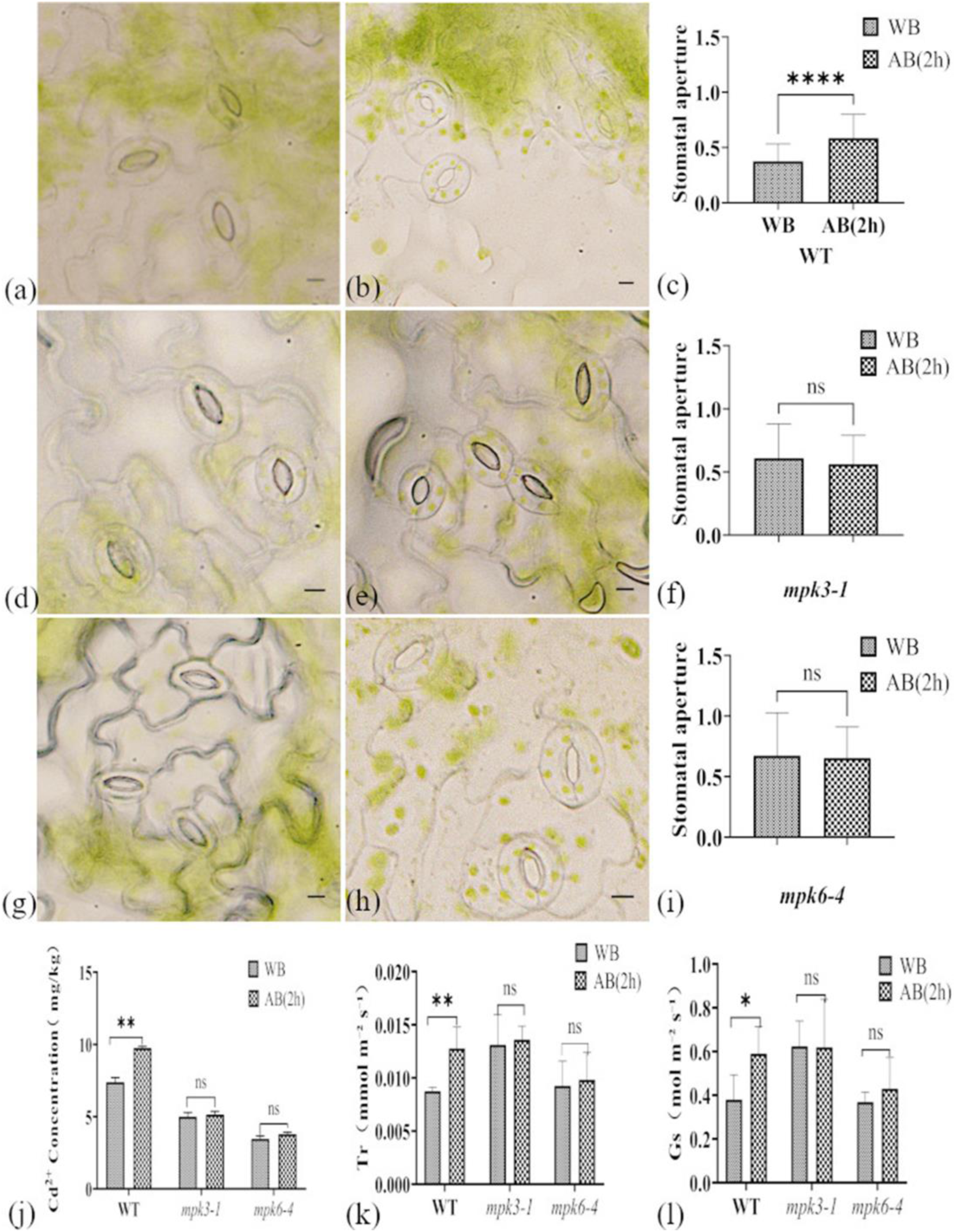
Trichome brushing does not affect stomatal aperture, Cd²⁺ accumulation, transpiration rate, or stomatal conductance in *mpk3-1* and *mpk6-4* mutants. Representative stomata in epidermal peels of WT (a, b), *mpk3-1* (d, e), and *mpk6-4* (g, h) leaves without (a, d, g) or with trichome brushing (b, e, h). Quantification of stomatal aperture in WT (c), *mpk3-1* (f), and *mpk6-4* (i) leaves. Cd^2+^ concentration in aboveground tissues (J), transpiration rates (k), and (l) stomatal conductance of WT, *mpk3-1* and *mpk6-4* plants. All plants were grown on Cd²⁺-supplemented MS medium. Scale bar: 40 µm. Statistical significance: ns, not significant (p > 0.05); *, p < 0.05; **, p < 0.01.

Because trichome brushing did not alter stomatal aperture in *mpk3-1* and *mpk6-4* mutants, we predicted it would likewise not affect Cd²⁺ accumulation, stomatal conductance, or transpiration rate. Consistent with this prediction, mechanical stimulation of trichomes did not significantly change shoot Cd²⁺ content, stomatal conductance, or transpiration rate in either mutant line (Fig. 6j–l; Fig. S9). In contrast, WT *Arabidopsis* exhibited pronounced increases in all three parameters following the same treatment.

Together, these results demonstrate that MPK3 and MPK6 are indispensable for trichome-stimulation-induced stomatal opening and enhanced Cd²⁺ uptake in *A. thaliana*.

### 4.6 Malic acid regulates stomatal opening and promotes Cd²⁺ accumulation following trichome stimulation

MPK3 and MPK6 have been reported to regulate stomatal immunity via malic acid metabolism (Su et al., 2017). Because malic acid promotes stomatal opening by balancing potassium fluxes in guard cells (Talbott and Zeiger, 1993; Santelia and Lawson, 2016), we hypothesized that trichome brushing alters malic acid levels in *A. thaliana*, thereby enhancing stomatal aperture and increasing shoot Cd²⁺ accumulation. qRT-PCR analysis revealed that trichome brushing significantly increased *MLS* transcript levels in WT plants, whereas brushing epidermal pavement cells reduced *MLS* expression in *gl1-1* mutants (Fig. 7a), indicating trichome-dependent regulation of malic acid biosynthesis. Consistently, malic acid content was elevated in WT leaves following mechanical stimulation, but decreased in *gl1-1* mutants (Fig. 7b).

**Fig. 7.**
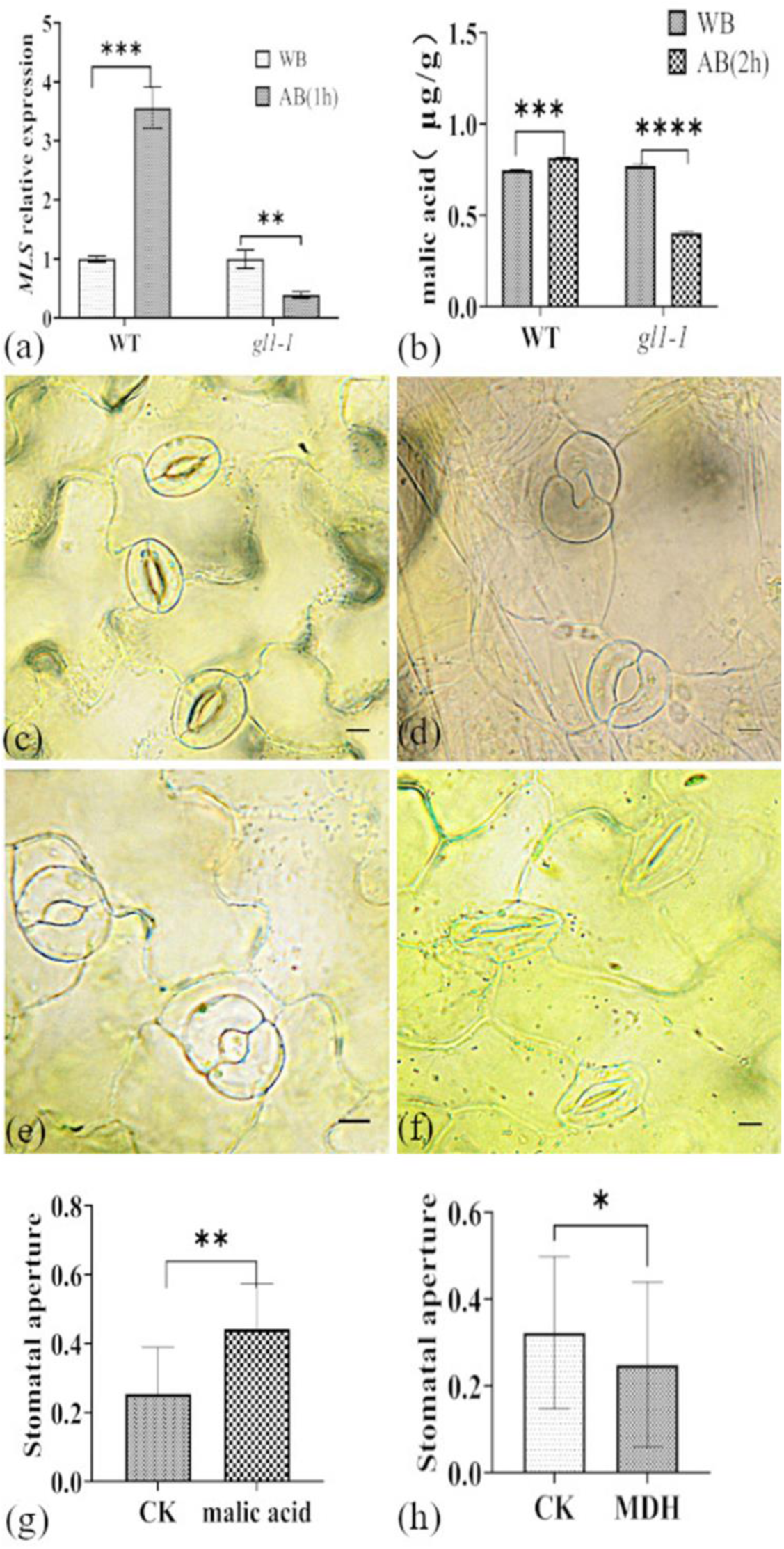
Trichome brushing alters *MLS* expression and malic acid levels, and exogenous treatments modulate stomatal aperture in WT leaves. (a) qRT-PCR analysis of *MLS* transcript levels and (b) malic acid concentration in WT and *gl1-1* leaves following trichome brushing. Representative stomata in epidermal peels of WT leaves under (c) untreated and (d) malic acid–treated conditions, and (e) before and (f) after MDH application. Quantification of stomatal aperture is shown in (g) and (h). All plants were grown on MS medium supplemented with Cd²⁺. Scale bar = 40 µm. Statistical significance: *, p < 0.05; **, p < 0.01;***, p < 0.001; ****, p<0.0001.

Pharmacological modulation further validated this mechanism. Exogenous malic acid (0.1 mM) significantly increased stomatal aperture in both WT and *gl1-1* leaves (Fig. 7c–d, g; Fig. S10a–c), whereas treatment with MDH (0.1 mM) reduced aperture in both genotypes (Fig. 7e–f, h; Fig. S10d–f). Moreover, malic acid application elevated Cd²⁺ levels in aerial tissues of both WT and *gl1-1* leaves (Fig. 8a–b), with a pronounced increase in Cd²⁺ accumulation specifically in WT trichomes (Fig. 8c–d).

**Fig. 8.**
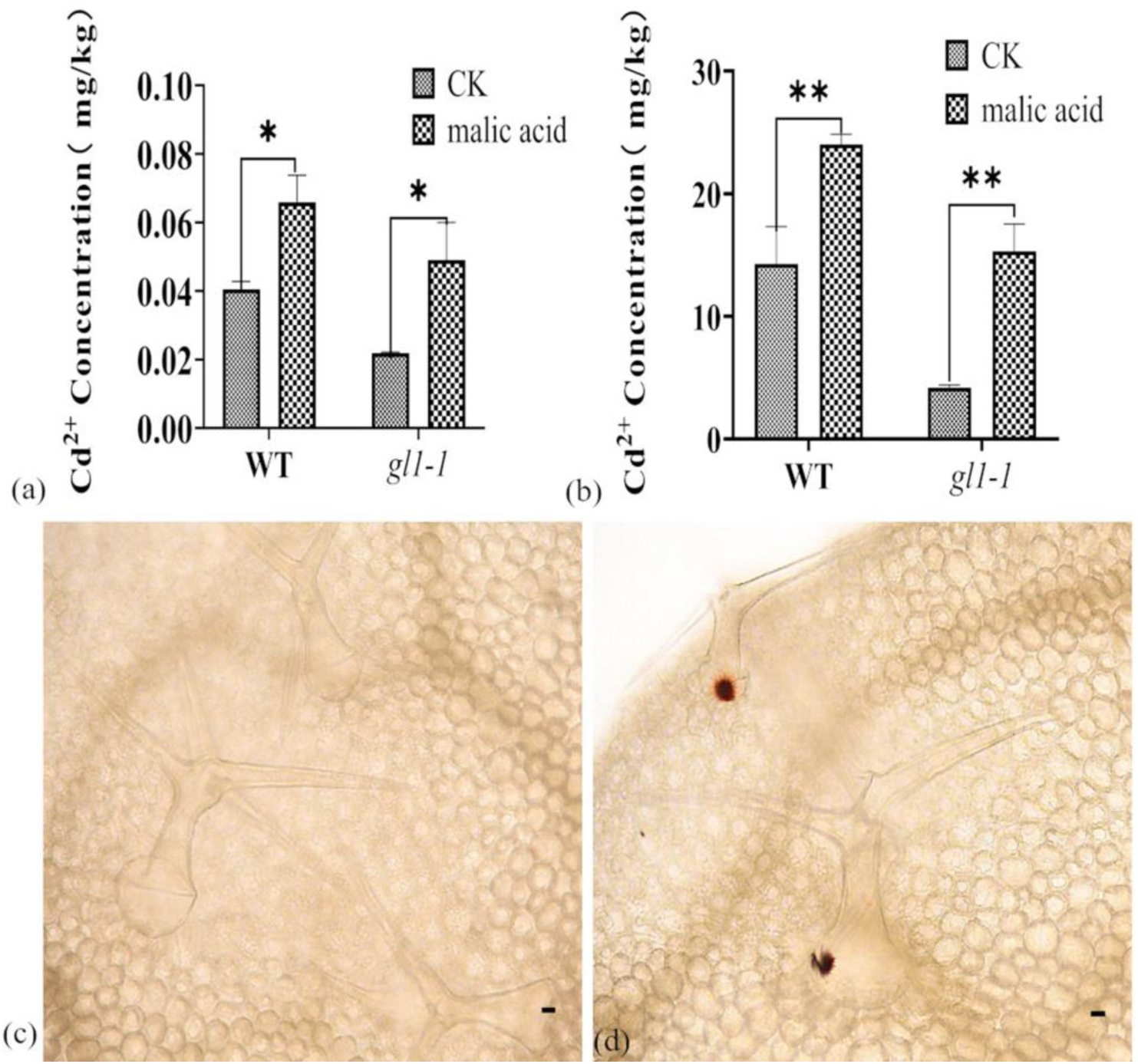
Exogenous malic acid enhances Cd²⁺ accumulation in *Arabidopsis* shoots and trichomes. (a) Shoot Cd²⁺ concentrations in WT and *gl1-1* plants under control conditions (no added Cd). (b) Shoot Cd²⁺ concentrations in WT and *gl1-1* plants following treatment with 5 µM CdCl₂. Data are mean ± SD. (c) Representative dithizone staining of Cd²⁺ in WT trichomes without malic acid. (d) Dithizone staining of Cd²⁺ in WT trichomes after incubation with 0.1 mM malic acid for 3 h. Scale bar = 40 µm. Statistical significance: *, p < 0.05; **, p < 0.01.

Together, these findings identify malic acid as a central regulator linking trichome-derived mechanical stimulation to stomatal aperture enlargement and Cd²⁺ accumulation.

## Discussion

This study establishes that mechanical stimulation of trichomes in *A. thaliana* initiates a signaling cascade that significantly alters leaf physiology. Trichome brushing increased stomatal aperture, conductance, and transpiration, responses absent in the trichome-deficient *gl1-1* mutant, confirming that trichomes are the primary mechanosensory structures. Elevated Cd²⁺ concentrations in aboveground tissues accompanied stomatal opening, linking mechanosensory input to ion accumulation.

Mechanistically, these responses were independent of ABA signaling and CAMTA3 but critically required MPK3 and MPK6. Loss of either kinase abolished stomatal opening, transpiration, and Cd²⁺ accumulation, correlating with reduced malic acid levels. Notably, MPK3/MPK6 activation during pathogen attack promotes stomatal closure by lowering malate (Su *et al*., 2017), whereas trichome stimulation produced the opposite outcome: increased *MLS* expression, elevated malate, and stomatal opening. This contrast underscores the context-dependent integration of upstream signals by the MAPK cascade, likely through differential regulation of malate metabolism or distinct downstream targets.

Although trichome stimulation raised ABA concentrations, stomatal aperture tracked malate rather than ABA, suggesting that malate overrides or antagonizes ABA’s canonical closing effect under mechanical stress. Exogenous malate enlarged stomata and enhanced Cd²⁺ accumulation, whereas MDH treatment suppressed stomata opening, confirming malate’s functional role in coordinating guard cell behavior and metal uptake. Future experiments should test whether *mpk3-1* or *mpk6-4* mutants fail to elevate malate following trichome stimulation, thereby directly establishing MAPK control of malate biosynthesis in this mechanosensory context. Beyond stomatal regulation, malate may also contribute to Cd²⁺ detoxification. Elevated leaf malate could chelate Cd²⁺ in the apoplast or cytosol, restrict its entry into mesophyll cells, or modulate intracellular compartmentalization. These possibilities are consistent with established functions of organic acids in mitigating heavy metal stress (Ma et al., 2001; Ma et al., 2020; Guan et al., 2024).

The mechanosensory response observed here may be broadly conserved among plants bearing non-glandular trichomes, which share structural susceptibility to mechanical stress. The non-glandular trichomes of *A. thaliana* exemplify a widespread class of slender epidermal structures found across diverse plant taxa (Watts and Kariyat, 2021). Mechanically, such slender structures are inherently prone to buckling or destabilization under external force (Ji et al., 2004). Given this shared vulnerability, we propose that the mechanosensory pathway leading to stomatal opening may represent a conserved adaptive response among plants with similar trichome architectures.

Finally, Cd²⁺ accumulation patterns differed between soil-grown plants under prolonged stress and plate-grown plants at earlier developmental stages, suggesting the developmental phase strongly influences Cd²⁺ partitioning. These divergent outcomes highlight the importance of growth context in interpreting trichome-mediated ion dynamics.

## Conclusion

Our research demonstrates that mechanical stimulation of trichomes on *A. thaliana* leaves significantly influences key physiological processes. Trichome stimulation increased stomatal aperture, leading to enhanced stomatal conductance and transpiration rates, and also elevated Cd²⁺ concentrations in aboveground tissues. These responses were strictly dependent on the presence of trichomes, as the trichome-deficient *gl1-1* mutant did not exhibit such changes. Mechanistically, we identified the MAPK pathway and increased malic acid levels as central signaling components mediating stomatal opening. We propose that this linkage between mechanical trichome stimulation and stomatal regulation may represent a conserved response among plants with non-glandular trichomes. Future work should clarify how the MAPK pathway governs malic acid accumulation and explore downstream genetic programs controlling Cd²⁺ transport and accumulation.

## Contribution

Li Hong Zhou conceived the study, analysed the data, and wrote the manuscript. Together, Lihong Zhou and Juan Du designed and conducted the study. Hongyang Wei, Tongxin Shen, Xue Chang contributed to the histochemical analysis, while Mingyu Hao and Yang Qi assisted with observing stomatal apertures. Zhiyan Cao, Ning Liu, and Jinguao Dong contributed to the discussion. All authors have read and approved the final manuscript.

## Acknowledgements

This work was supported by the Natural Science Foundation Hebei Province (No.202204052) and the Project for the Central Guided Local Science and Technology Development Fund (No.236Z2901G). We appreciate Prof. Yu Diqiu from Yunnan University, Prof. Shuhua Yang from China Agricultural University, and Associate Prof. Houqing Zeng from Hangzhou Normal University for providing us with *gl1-1*, *mpk3-1/mpk6-4*, and *sr1-2*/*CAMTA3*-*ox5* seeds.

